# Damped White Noise Diffusion with Memory for Diffusing Microprobes in Ageing Fibrin Gels

**DOI:** 10.1101/576710

**Authors:** R.R.L. Aure, C.C. Bernido, M.V. Carpio-Bernido, R.G. Bacabac

## Abstract

From observations of colloidal tracer particles in fibrin undergoing gelation, we introduce an analytical framework that allows determination of the probability density function (PDF) for a stochastic process beyond fractional Brownian motion. Using passive microrheology via videomicroscopy, mean square displacements (MSD) of tracer particles suspended in fibrin at different ageing times are obtained. The anomalous diffusion is then described by a damped white noise process with memory, with analytical results closely matching experimental plots of MSD and PDF. We further show that the white noise functional stochastic approach applied to passive microrheology reveals the existence of a gelation parameter *μ* which elucidates the dynamics of constrained tracer particles embedded in a time dependent soft material. This study offers experimental insights on the ageing of fibrin gels while presenting a white noise functional stochastic approach that could be applied to other systems exhibiting non-Markovian diffusive behavior.

## INTRODUCTION

Fibrin is a protein polymer network that plays a crucial role in blood clotting, wound healing, and other biological processes such as blood vessel formation and cell growth (1–6). Aside from its importance in clinical applications, fibrin also finds an increasing role in tissue regeneration and engineering (5, 7–8). Thus, it is important to have a quantitative description of the mechanics and complex structural formation of fibrin as a viscoelastic biomaterial (3, 9–11). Since not much is known about the diffusive behavior of probe particles in fibrin as its structure evolves in time, we use passive microrheology via video microscopy to reveal the gelation dynamics of fibrin by tracking the movement of probe particles suspended in its network of protein polymers (12–14). Moreover, to elucidate how the structure of fibrin is affected at different ageing times, we present a theoretical framework for the stochastic fluctuations exhibited by the experimental observations.

The diffusion of colloidal tracer particles in complex fluids is usually analyzed by measuring the mean square displacement (MSD), < Δ*x*^2^(*τ*) >, where brackets represent an ensemble average and *τ* is the lag time between two positions taken by the particle in its trajectory. Numerous studies have interestingly revealed anomalous diffusion or deviations from purely Brownian motion (15–26) thus making studies of probe particles in many biophysical systems an active area of research. Examples of systems where anomalous diffusion occurs include entangled F-Actin networks (17–19), living cells (20, 21), cytoplasm (22, 23), colloidal liquids (24, 25), and lipid membranes (26), among others. Anomalous diffusion, is often characterized analytically using fractional Brownian motion (fBm), which follows power law scaling of the form (13, 15), < Δ*x*^2^(*τ*) > ~ *τ^α^*. In particular, the motion of the probe particles suspended in the sample is said to be subdiffusive when *α* < 1, and superdiffusive when *α* > 1. Ordinary Brownian motion is recovered when *α* = 1. However, as shown by our experimental results, there is a need to go beyond fBm for a suitable predictive mathematical model that can closely reflect the empirically obtained MSD which may not necessarily follow pure power law scaling. Here we show how anomalous diffusion is described by a damped white noise process with memory, with analytical results more closely matching experimental plots of MSD and PDF. This paper then aims to contribute useful information on the underlying dynamics of fibrin gelation and network formation while presenting a white noise functional stochastic approach that could be applied to other systems exhibiting non-Markovian diffusive behavior.

## MATERIALS AND METHODS

### Sample Preparation

Human fibrinogen (depleted of plasminogen, von Willebrand factor, and fibronectin) and α-thrombin were obtained in powder form from Enzyme Research Laboratories (South Bend, IN). The fibrinogen solutions were prepared in buffer solutions of pH 7.4 containing 200 mM HEPES, 1500 mM NaCl, 5 mM CaCl_2_, and Milli-Q water. The resulting solution was mixed using a vortex mixer for ~ 5 s. The protein concentration of 3.0 mg/ml was prepared for this experiment. Pre-diluted polystyrene microspheres with diameter of 1 μm purchased from Phosphorex, Inc. (Hopkinton, MA) were added into the mixtures. To break up the aggregates of micron sized microspheres and produce a homogeneous mixture, the resulting solution was agitated with a sonicator for 10 minutes and mixed vigorously for ~ 30 seconds with a vortex mixer. To initialize polymerization, the fibrinogen solutions were added to 0.5 U/ml thrombin. Finally, the sample was pipetted into a sample chamber consisting of two adhesive spacers between a glass microscope slide and a cover slide and sealed with grease to prevent drift.

### Particle Tracking

To observe how the motion of the colloidal tracer particles is affected by the formation of fibrin networks as the sample ages, we transferred the sealed sample chamber to the optical microscope for imaging. The particle tracking experiments were performed on a bright field optical microscope using an oil immersion 60×, 1.25 NA objective lens (see Movie S1 and Movie S2 in the Supporting Material). To minimize the effect of unwanted heating which might affect the properties of the sample due to the light source, a specialized green filter disc was placed in between the light beam source and the sample. Furthermore, the images were scanned at the middle portion of the glass slide and cover slip interface in order to minimize interactions with the cover slip. Videos were recorded with a CCD camera (Watec WAT-902H) at a frame rate of 25 frames per second for one minute (1500 frames) at different ageing times. The ageing time, t_1_, is defined to be the time interval starting from the addition of thrombin to recording each video which is a slight modification of the definition of Houghton et al. (27). The movies of the changing positions of the diffusing colloidal probe particles were then analyzed using Trackpy v0.4.1 which is a Python package for particle tracking and a full description can be found online (28). Based on the trajectories of the colloidal tracer particles suspended in the sample we calculated the mean square displacement at different ageing times (t_1_ = 6 min, t_2_ = 15 min, t_3_ = 20 min, t_4_ = 25 min, t_5_ = 35 min, t_6_ = 40 min, and t_7_ = 60 min). All measurements were performed at a room temperature of 20°C and the experiment was repeated four times using a fresh sample of reconstituted fibrin.

### Theoretical Framework

To provide insights on the anomalous diffusion of tracer particles embedded in fibrin, we introduce a damped white noise process with memory for analyzing the modes of diffusive behavior valid for different gelation stages or ageing times. Guided by insights from the use of different memory kernels for stochastic processes in biological systems (29, 30), we parametrize the fluctuating position *x*(*τ*) of the tracer particle as,

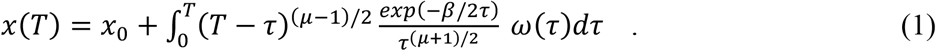

Here, *ω*(*τ*) is the white noise random variable, i.e., *ω*(*τ*) = *dB*(*τ*)/*dτ*, with *β*(*τ*) an ordinary Brownian motion (31). The factor, *exp*(–*β*/2*τ*)/*τ*^(*μ*+1)/2^, dampens the white noise variable, while (*T* – *τ*)^(*μ*-1)/2^ serves as a memory function as diffusion starts at *x*_0_ at time *τ* = 0 up to the current time *τ* = *T*. Parameters *μ* and *β* have adjustable values depending on the system being studied.

As parametrized by Eq. 1, the fluctuating position of the probe particle can be anywhere at some later time *T*. To satisfy the boundary condition of observed position *x*(*T*) as the point *x_T_*, the Donsker delta function (31) constraint is applied,

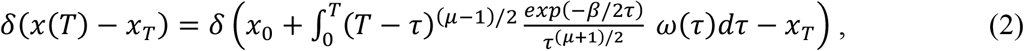

where we have written *x*(*T*) explicitly using Eq. 1. For all fluctuating paths that do not terminate at *x_T_*, the delta function Eq. 2 vanishes. One can then obtain the probability density function, *P*(*x_T_,T;x*_0_,0), that a path ends at *x_T_* if it started at *x*_0_, by integrating over all possible paths that satisfy Eq. 2, i.e.,

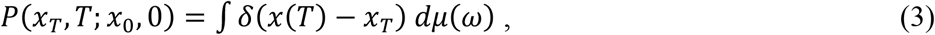

where, 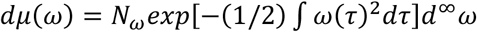, is the Gaussian white noise measure (31). The *N_ω_* is a normalization factor and the exponential is responsible for the Gaussian fall-off. Expressing the delta function in terms of its Fourier representation we have,

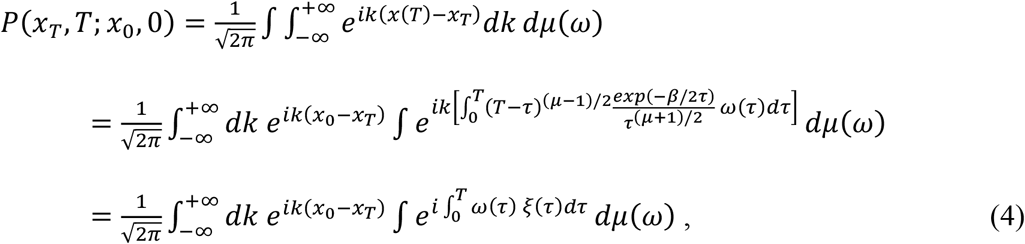

where we let, 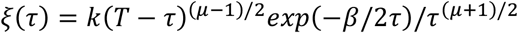. Integration over *dμ*(*ω*) is done using the characteristic functional (31),

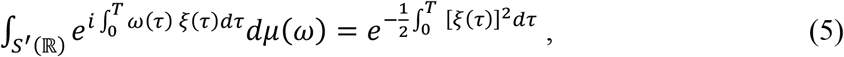

with, *ξ* ∈ *S*(ℝ), where *S*(ℝ) and *S*′(ℝ) are the real Schwartz spaces of test functions and tempered distributions, respectively. The Gel’fand triple, *S*(ℝ) ⊂ *L*^2^(ℝ) ⊂ *S*’(ℝ), links the space of test functions *S*(ℝ) with *S*′(ℝ) through the Hilbert space of square integrable functions *L*^2^(ℝ). With Eq. 5 we get from Eq. 4,

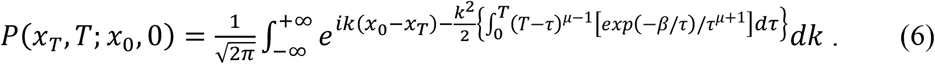

The integral over *dk* is a Gaussian integral which yields a probability density function (PDF) *P*(*x_τ_, T; x*_0_,0) given by,

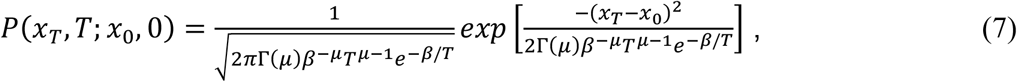

where we used Eq. 3.471.3 of reference (32) for integration over *dτ*. Here, Γ(*μ*) is the gamma function, and *x_T_* is the position of a diffusing tracer particle at time *T*. As can be shown, this PDF satisfies a modified diffusion equation of the form (30, 33),

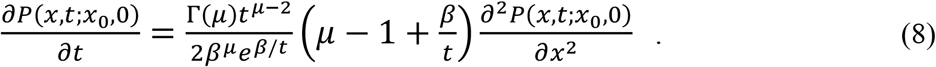

For a stochastic process described by Eq. 7 an evaluation of the mean square displacement, MSD = < Δ*x*^2^(*τ*) >, yields (30),

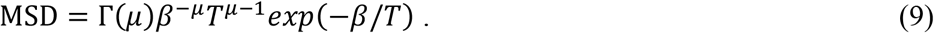

With Eqs. 7 and 9, we are able to characterize the gelation process with a close match between a nonlinear empirical and theoretical MSD obtained as shown in the Results section. Moreover, *μ* turns out to be a gelation parameter which could track the amount of gelation that has transpired.

## RESULTS AND DISCUSSION

To obtain insights into the structure of fibrin networks as they evolve over time, we used passive particle tracking of a fluctuating probe particle at different ageing times (from t_1_ = 6 min to t_7_ = 60 min) defined as the time interval starting from the addition of thrombin to the start of each video recording. A sample trajectory of a probe particle embedded in fibrin taken 20 minutes after the addition of thrombin is shown in Figure 1.

**Figure 1.**
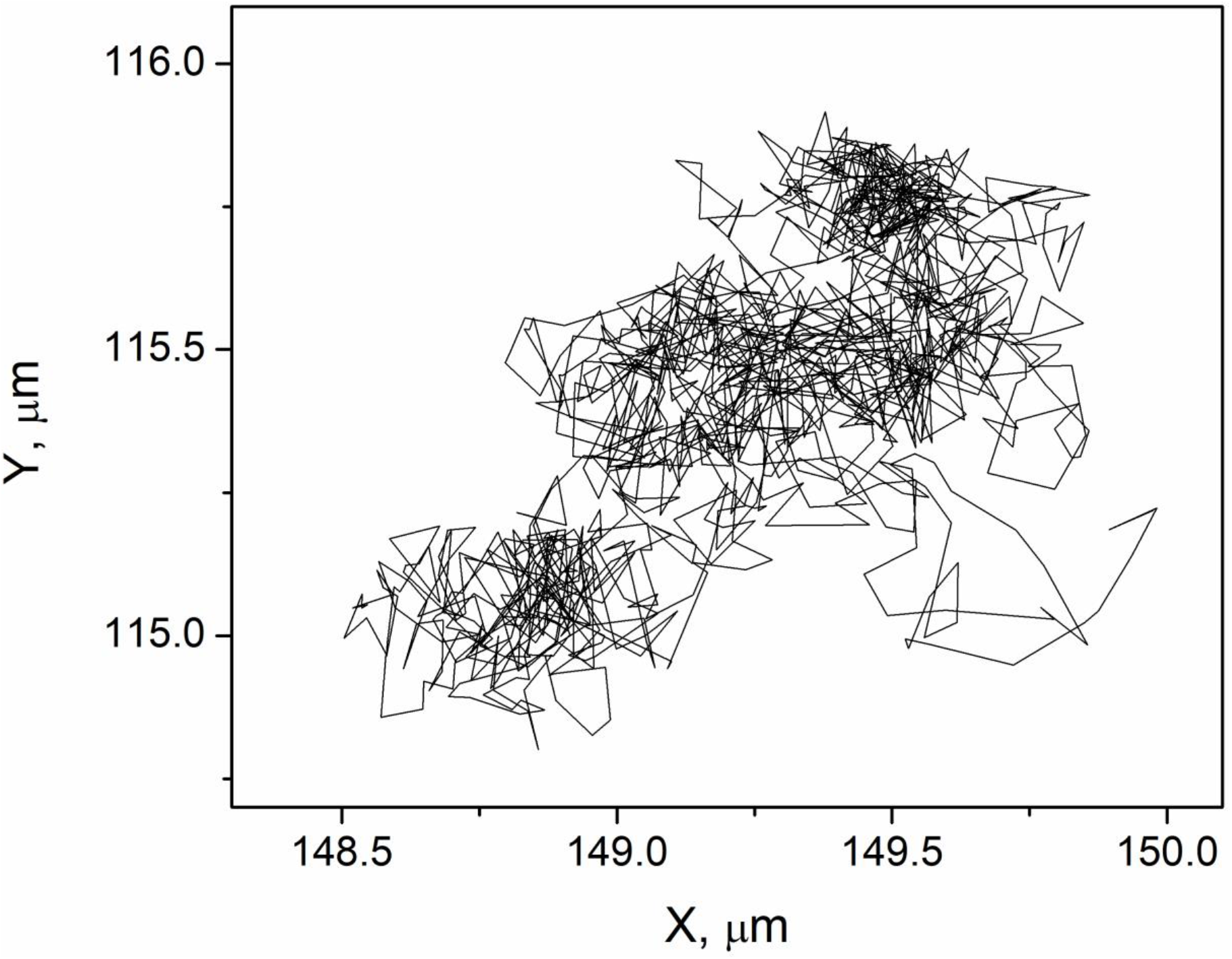
Sample trajectory of a tracer particle embedded in fibrin at ageing time, *t*_3_ = 20 min, from video microscopy.

For fluctuations of tracer particles along one dimension, the mean square displacement (MSD) is logarithmically plotted in Figure 2 for MSD as a function of the lag time *τ* at different ageing times. Noticeable systematic transition from purely diffusive motion to subdiffusive behavior is seen from the graphs of the MSD’s. As the ageing time increases, the MSD exhibits different diffusive behaviors at the short, intermediate, and longer lag times *τ*. These results support the findings of earlier works on fibrin exhibiting a hierarchical and complex structure (3, 9, 34). The fibrin network clearly causes the tracer particles to undergo anomalous diffusion as the sample ages.

**Figure 2.**
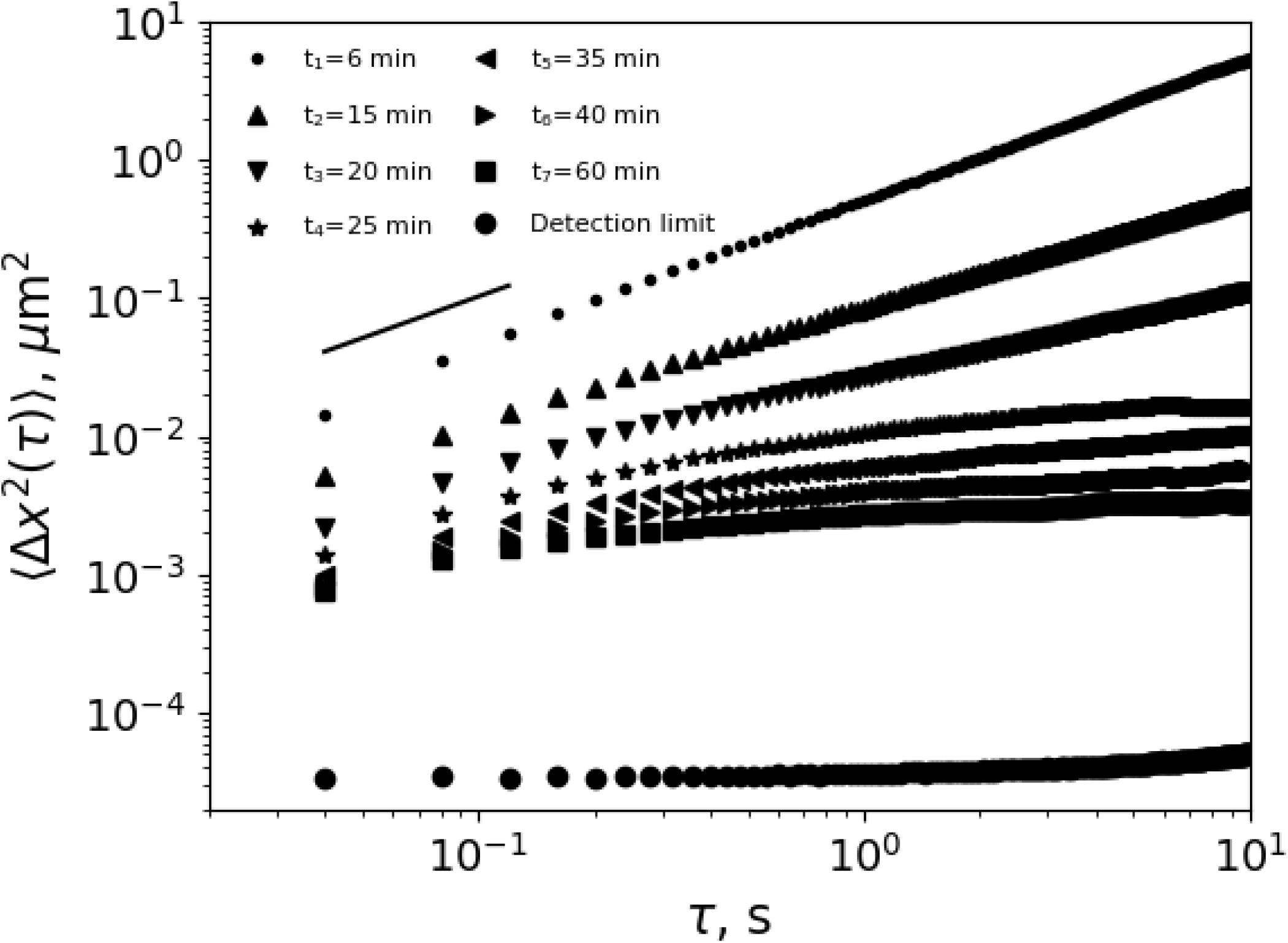
Plot of one-dimensional empirical MSD versus lag time of the tracer particles embedded in fibrin at different ageing times. The solid line indicates a linear fit.

As shown in Figure 2, at early ageing time, t_1_ = 6 min, the MSD curves of the mobile particles behave linearly with lag time τ showing that the sample is still liquid-like. However, as the ageing time increases to t_*a*_ ≥ 15 min, the plot of the MSD significantly deviates from the linear dependence to lag time. Noticeable features include the shift in the slope of the MSD as lag time *τ* increases and the onset of a plateau visible when the ageing time t_α_ > 25 min. This feature in the MSD is reported as an indication that the tracer particles in the sample are constrained or trapped by the surrounding network structures (27, 35–36). Note that the measured MSD’s at different ageing times are well above the detection limit of 3 × 10^−5^*μ*m^2^ determined by measuring the apparent MSD of immobile probe particles in the sample chamber.

Let us now analytically address the underlying mechanism of the anomalous diffusion of probe particles when the fibrin networks evolve in time. We start by noting that if we use fractional Brownian motion (fBm) with an MSD of the form, < Δ*x*^2^(*τ*) > ~ *τ^α^*, its log-log plot is always a straight line which fails overall to capture a curved empirical MSD (see Fig. S1 in the Supporting Material). With fBm, as seen in Figure 3, one needs to change from a value of *α* ≈ 0.9 to *α* ≈ 0.6 to match an empirical MSD as one goes from shorter to longer lag times *τ*. This change to a lower value of *α* at longer lag time scales, however, indicates that another process is observed that possibly arises due to the presence of network structures in the sample constraining the motion of the embedded tracer particles.

**Figure 3.**
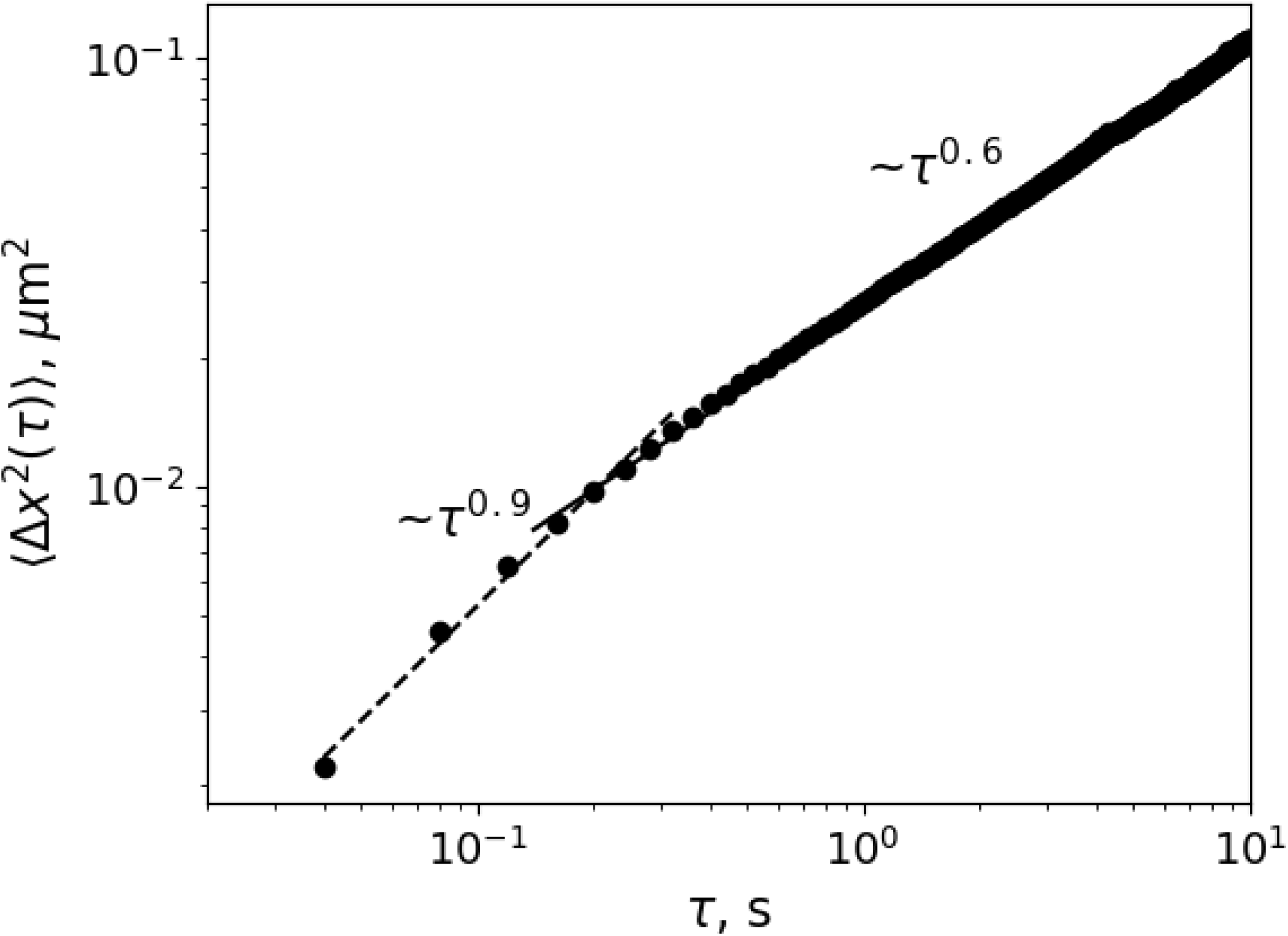
Empirical MSD (circles) for an ageing time of 20 minutes. For fractional Brownian motion with < Δ*x*^2^(*τ*) > ~ *τ^α^* to match a curved MSD, two parameter values are used: *α* ≈ 0.9 (shorter lag times; dash line) and *α* ≈ 0.6 (longer lag times; solid line).

Going beyond fBm, however, a damped white noise process with memory described by Eqs. 7 to 9, more naturally captures the behavior of tracer particles for different ageing times as shown in Figure 4. For each ageing time, a good match for the empirical and theoretical MSD’s is obtained with fixed values of *β* and *μ* as one goes from shorter to longer lag times *τ*, unlike that of fBm where *α* has to change (see, e.g., Fig. 3). Moreover, across different ageing times, the same value of *β* = 0.05 s is used, while parameter *μ* decreases in value as the sample ages (see Table 1 and Figure 5). This suggests that *μ* may be identified as an ageing or gelation parameter for fibrin networks as time progresses. The theoretical graphs of Fig. 4 (dash lines) are normalized by multiplying Eq. 9 by a constant *c*, i.e., *c*(MSD). The constant *c* raises or lowers the theoretical MSD plot without changing its shape so that a good overlap with the empirical MSD is obtained for easy comparison. Note, however, that the first value of *τ* in Figure 4, typical of the start of video recording, shows a slight discrepancy between theory and experiment. Furthermore, the experiments were also repeated for different fibrinogen concentrations (see Fig. S2 in the Supporting Material). It can be seen that reasonable agreement between the empirical MSDs and theory for different fibrinogen concentrations is observed suggesting that indeed the damped white noise process with memory can robustly describe the dynamics of the embedded tracer particles constrained by network structure in the sample. Furthermore, the experiments were also repeated for different fibrinogen concentrations and the damped white noise process with memory seems to capture the overall behavior of the empirical MSDs (see Fig. S2 in the Supporting Material).

**Table 1.** A close match between empirical and theoretical MSD is obtained for parameter values of *μ* corresponding to different ageing times. For the theoretical MSD, Eq. 9, *β* = 0.05 s for all values of *μ*.

**Figure 4.**
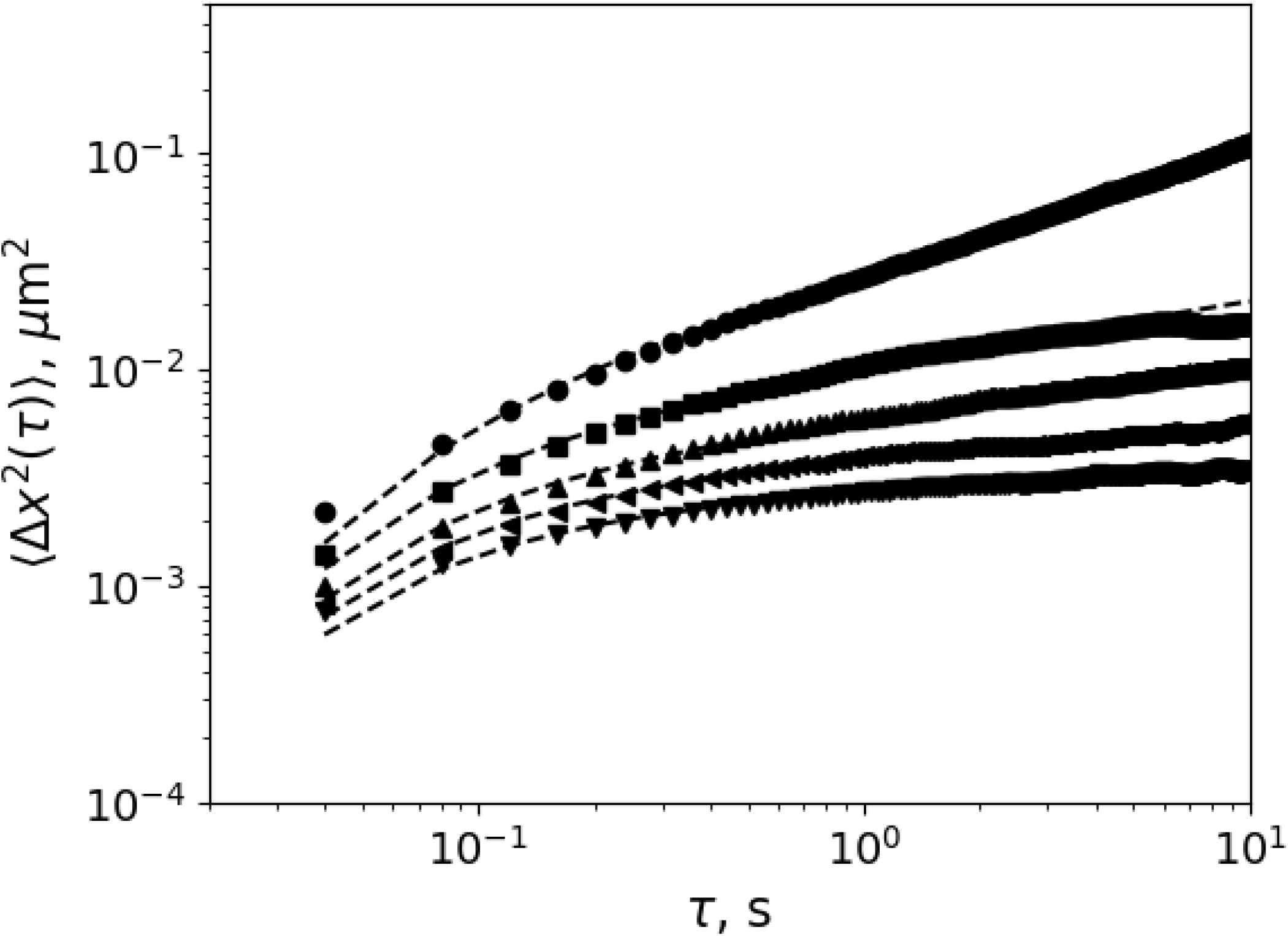
Plot of one-dimensional empirical MSD versus lag time of tracer particles embedded in fibrin at ageing times of 20 minutes (circle), 25 minutes (square), 35 minutes (triangle marker up), 40 minutes (triangle marker left), and 60 minutes (triangle marker down). The empirical MSD is matched with a normalized theoretical fit (dash lines, Eq. 9), with *β* = 0.05 s for different ageing times and decreasing values of *μ* as gelation progresses as shown in Table 1.

**Figure 5.**
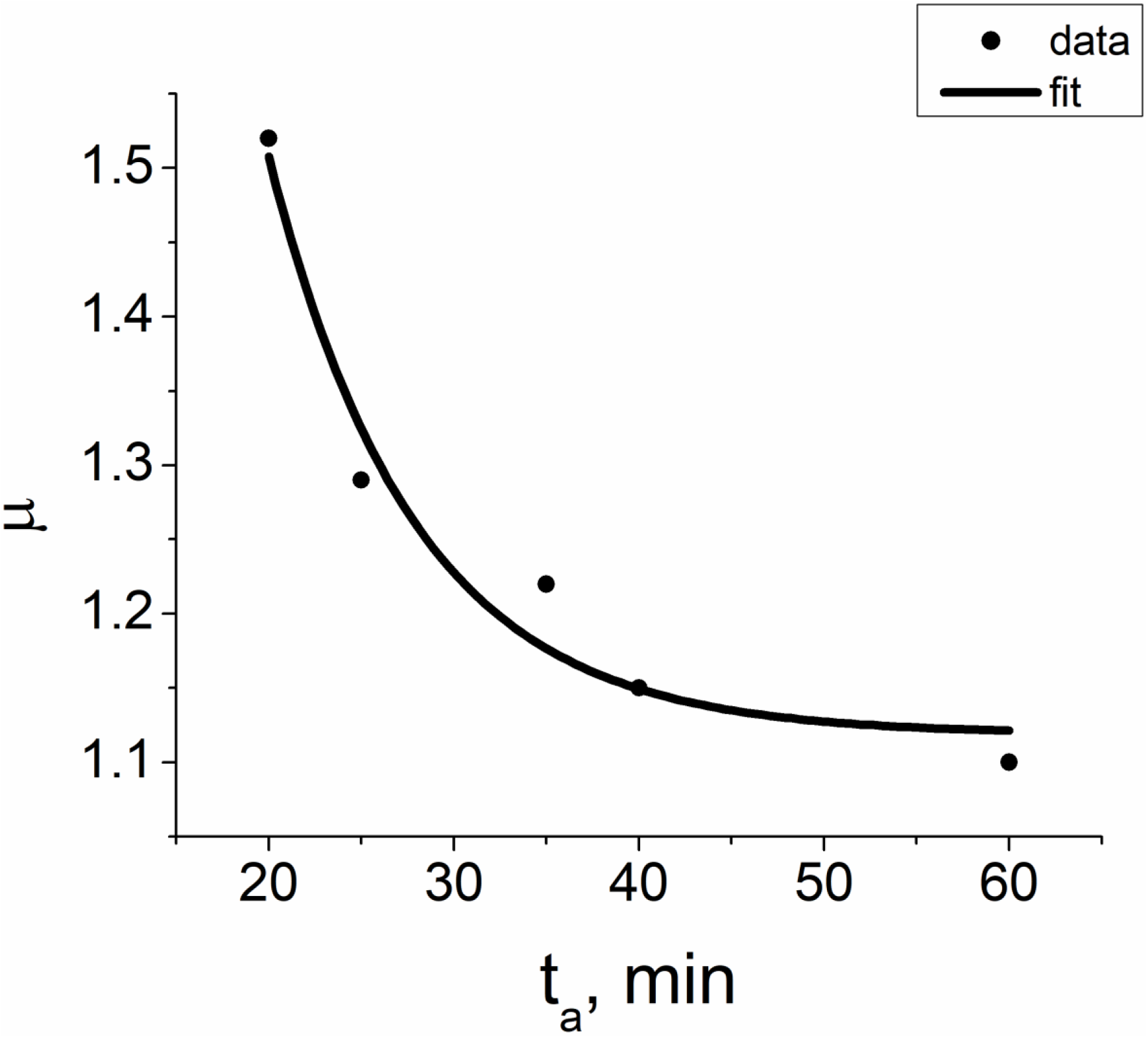
Plot of parameter *μ* as a function of ageing time. The solid curve represents an exponential fit of the data.

To have a better insight on the role of parameter *μ*, we can expand the exponential in Eq. 9 and write the MSD alternatively as,

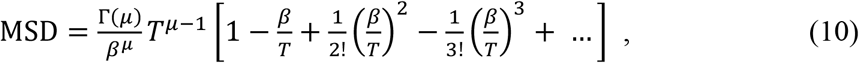

where time *T* corresponds to the empirical lag time *τ* in the particle tracking video. Clearly, Eq. 10 goes beyond fractional Brownian motion which has an MSD, < Δ*x*^2^(*τ*) > ~ *τ^α^*, commonly employed in studies of particle tracking experiments. However, if we let *μ* – 1 = *α*, we can view Eq. 10 as an fBm-like MSD with time-dependent corrections given by terms inside the square brackets. Note that higher order correction terms become negligible with *β* = 0.05 s. From short to longer lag times, these correction terms inside the square bracket enable the theoretical MSD to capture a curved log-log plot of the empirical MSD (Fig. 4) even if only one value of *α*, or *μ*, is used for each ageing time (see, Table 1), unlike that shown in Fig. 3 for fBm.

Given an ageing time, one can also plot the experimental distribution of displacements Δ*x* corresponding to a lag time, for example, *τ* = 0.04 s. This can then be compared with the theoretical probability density function, Eq. 7, where we have, Δ*x* = *x_T_* – *x*_0_, and *T* = 0.04 s. This comparison is shown in Figure 6 for ageing time, t = 40 min. Other ageing times give similar results where a close match is obtained between empirical and theoretical plots.

**Figure 6.**
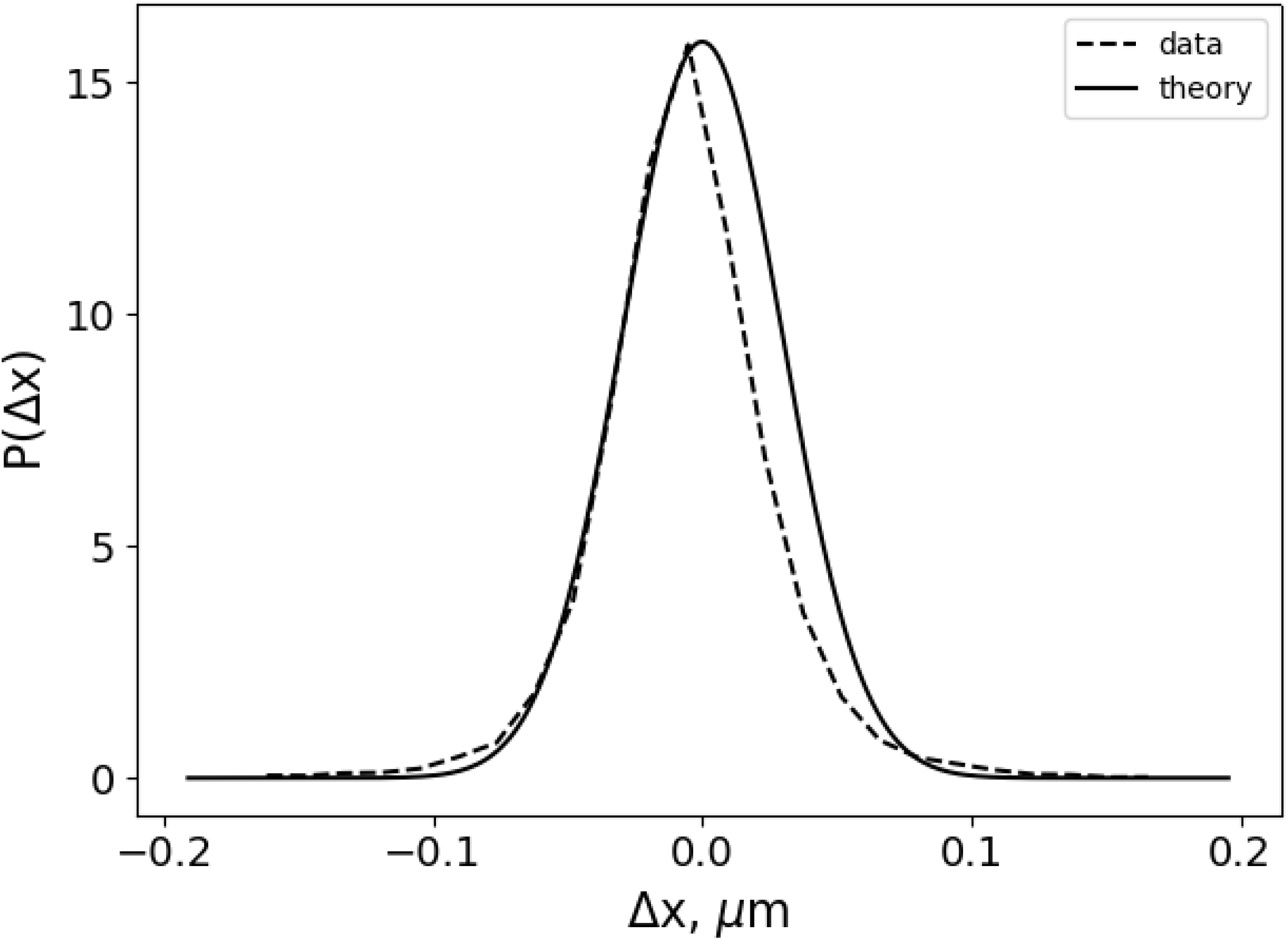
Experimental distribution of displacements Δ*x* (dash line) for lag time *τ* = 0.04 s and ageing time t = 40 min. Theoretical fit (solid line), Eq. 7, for *T* = 0.04 s, *μ* = 1.15 and *β* = 0.05 s.

One could also track the behavior of the probability distributions for Δ*x* by plotting them as ageing time increases as shown in Figure 7. For both the experimental and theoretical (dashed and solid lines) plots, the range of Δ*x* decreases as ageing time increases with a corresponding rise in the peak of the PDF around Δ*x* = 0. This implies a more constrained motion of the probe particles as gelation time increases. Note that for the theoretical PDF plot in Figure 7, the values *β* = 0.05 s and *T* = 0.04 s are used in Eq. 7 for different ageing times, and only the value of parameter *μ* is changed as given by Table 1.

**Figure 7.**
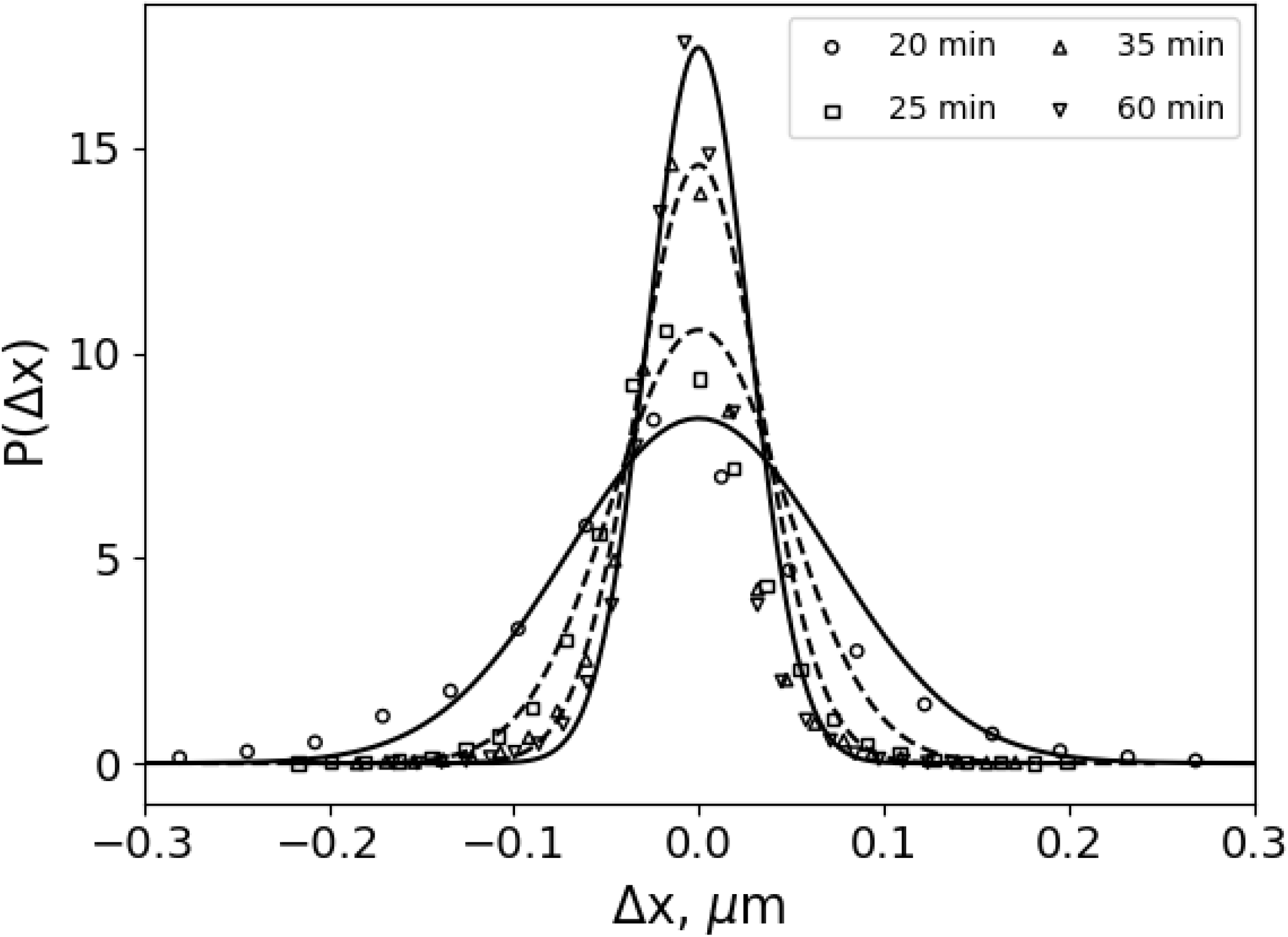
Probability distribution of displacements Δ*x* for lag time *τ* = 0.04 s at different ageing times. Experimental (symbols); Theoretical, Eq. 7 (dashed and solid lines).

## CONCLUSION

In view of its wide-ranging use from clinical applications to tissue engineering, it is important to investigate the ageing property of fibrin. For this, we use passive microrheology via videomicroscopy to investigate fluctuating motions of embedded tracer particles in fibrin at different time intervals from 6 minutes up to 60 minutes. Empirical mean square displacement (MSD) of the probe particles are then plotted for different ageing times. An analytical description of the observed fluctuations is facilitated by introducing a damped white noise process with memory, Eqs. 7 to 9, which allows a good match between theoretical and experimental MSD and PDF. Moreover, a parameter *μ* is identified as an ageing or gelation parameter whose value decreases as time increases. The probability density function of the novel non-Markovian stochastic process satisfies a modified diffusion equation which could provide additional insights for the analysis of gelation properties of fibrin as well as provide a functional stochastic approach that could be applied to other systems exhibiting non-Markovian diffusive behavior beyond fractional Brownian motion.

## AUTHOR CONTRIBUTIONS

R. R. L. A. and R. G. B. designed and performed the experiment. C. C. B. and M. V. C.-B. contributed analytical tools. R. R. L. A., C. C. B., M. V. C.-B., and R. G. B. analyzed the data. R. R. L. A., C. C. B., and M. V. C.-B. wrote the manuscript.

## ACKNOWLEDGMENTS

Useful discussions with the USC Medical Biophysics Group are gratefully acknowledged. R. R. L. A. wishes to thank the Commission on Higher Education-FDP-II and Visayas State University for financial support. R. G. B. was funded by the Philippine Council for Industry, Engineering and Emerging Technology Research and Development-Department of Science and Technology project no. 04310 and received logistic support from the USC Research Office and Department of Physics.

